# Uncovering Features of ProteinMPNN with Sparse Autoencoders

**DOI:** 10.64898/2026.11.09.765161

**Authors:** Neel Mukkavilli, Konstantin Ivanov, David C. Anastasiu, David A.C. Beck, Hamed Akbari, Michelle E. McCully

## Abstract

Customization and modification of machine learning derived sequences can be achieved by understanding the mechanics behind state-of-the-art protein design models such as ProteinMPNN. Uninterpretability is hypothesized to be due largely to polysemanticity where neural networks store more features than they have dimensions. Here, we introduce sparse autoencoders (SAEs) into ProteinMPNN which expand its latent space and uncover correlations between the activations in sparse dimensions and the presence of protein features such as secondary structure, residue volume, residue shape, disulfide bridges, and coordination to ligands. Additionally, we show that sparse activations can be manipulated to produce a variant of the short chain variable fragment Brolucizumab that is more thermostable than variants produced by traditional machine learning methods based on high temperature molecular dynamics simulations. Thus, SAEs provide both mechanistic understanding of protein design models and allow customizable manipulation of sequence prediction.

## 1. Introduction

Protein language models have transformed the landscape of protein science, enabling models to predict structure and sequence with levels of accuracy that rival traditional biophysical models Lin et al. (2023); Zhang et al. (2024); Ruffolo and Madani (2024); Madani et al. (2023); Chen et al. (2025); Komp et al. (2025). However, these models are unable to take into account the needs of the user who may require protein features at a given position such as a helix, catalytic site, hydrophobic pocket, etc. To remedy this, the geometric and energetic relationships that protein design models already learn can be leveraged.

Message Passing Neural Networks (MPNNs) are a class of protein structure-to-sequence prediction models that aim to mimic physical models Ingraham et al. (2019). By treating proteins as spatial graphs, MPNNs embed atoms or residues as nodes connected by geometrically meaningful edges. These networks iteratively exchange information between nodes through “messages”, allowing them to capture local and global structural relationships. All this information is stored in a set of intermediate vectors known as the latent space. Dimensions in these vectors are often polysemantic, meaning they store multiple features to save space which creates noise and opaque mechanics Elhage et al. (2022).

By eliminating polysemanticity, the information the model learns about the protein becomes open to understanding and manipulation. We propose an interpretable framework for protein design that combines ProteinMPNN Dauparas et al. (2022)with Sparse AutoEncoders (SAEs). SAEs are a simple neural network that expands the dense latent space into an overcomplete and sparse latent space, so each feature is represented in its own dimension. SAEs have been shown to disentangle high-dimensional latent spaces into sparse, human-interpretable features in large language models Bills et al. (2023); Cunningham et al. (2024); Wang et al. (2022); Elhage et al. (2022); Bricken et al. (2023) and similarly in protein language models like ESMFold Simon and Zou (2025).

We aim to probe the sparse latent space of SAEs in ProteinMPNN to find which dimensions correspond to understood and measurable protein properties. This may answer open questions regarding which biophysical principles, such as residue environments, secondary structures, residue chemistry, or long-range interactions, guide sequence predictions.

Furthermore, the sparse latent space could be manipulated to increase the thermostability of short chain variable fragments (scFvs) outlined in Fig. 1. scFv proteins are highly specific therapeutics but limited by their stability and have been the focus of previous machine learning and de novo engineering attempts Jamali et al. (2025); Akbarian and Chen (2022); Tadokoro et al. (2024); Wörn and Plückthun (2001); Harmalkar et al. (2023); Kalantar et al. (2024); Tu et al. (2016). By combining knowledge of protein mechanics with the control of interpretable features and the model’s understanding, SAEs may allow more effective, customizable, and robust thermostablization and potentially introduce transparency and reliability into machine learning driven protein engineering more broadly.

**Figure 1.**
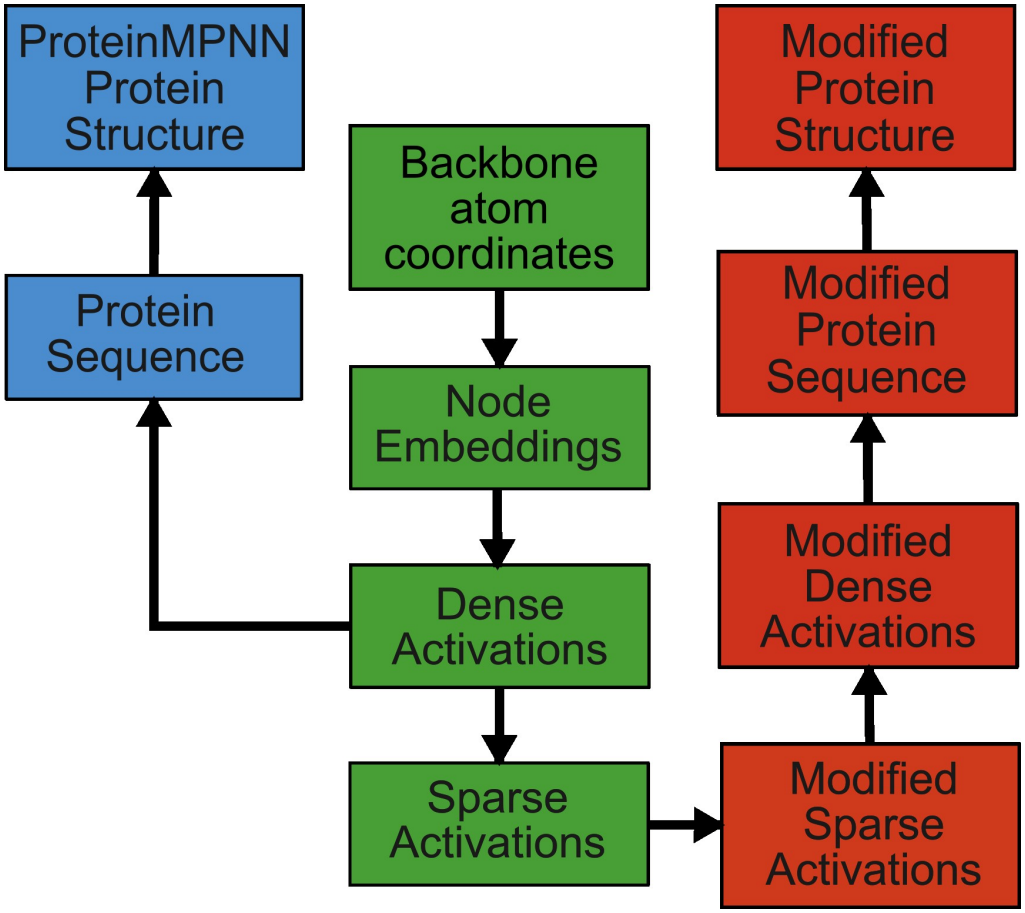
Diagram of relationship between native, ProteinMPNN, and modified protein variants used for steering

## 2. Methods

### 2.1. Sparse Autoencoders

ProteinMPNN is composed of an encoder and decoder. The encoder creates the final latent representation, and the decoder uses every residue’s representation to predict the amino acid identity. This work focuses on the encoder since, conceptually, the output of the encoder should be the most information-rich representation of the protein.

The sum of information between all of a node’s relevant edges, the message, updates the node’s latent space. The message for every edge’s corresponding nodes then updates the edge’s latent space. This repeats three times for three encoder layers. After each layer, an SAE used the dense latent space as its input, expanded it into a sparse latent space, and encoded it back. Only the node latent space is evaluated since it contains residue-specific information that is easier to evaluate. While some additional information may be in the edge latent space, it is much larger and therefore more computationally expensive to evaluate.

#### 2.1.1. Training Dataset

The original ProteinMPNN code repository was cloned, and the trained ProteinMPNN weights were frozen and used for training. The same dataset that was used for training ProteinMPNN was used for training the SAEs.

During SAE training, training and evaluation clusters were sampled in the same way as ProteinMPNN, but structure loading, batch size, and cluster size, were modified to accommodate the system RAM limits (32 GB). The training data was used as input for the ProteinMPNN encoder, and the latent space generated became the training data used for each SAE. SAE encodings were not fed back into the model, so each SAE was trained independently. There was no overlap between the training and evaluation datasets.

#### 2.1.2. Architecture and Loss Function

All ProteinMPNN node and edge embeddings were represented as a 128-dimensional vector. Thus, each SAE received and output a 128-dimensional vector (a full node array of size [# of batches, # of residues, 128]. Four different SAEs were trained to expand the vector to 1, 2, 4, and 8 times the original size in all three layers (message passing steps) of the encoder (referred to as SAE128, SAE256, SAE512, and SAE1024). We chose to train L1 penalty SAEs based on Elhage et al. (2022); Bricken et al. (2023); Cunningham et al. (2024); Simon and Zou (2025). The SAEs used a ReLU activation function and pre-encoder biases were applied and fixed to the bias of the decoder. Thus, the forward pass for an SAE was defined as:

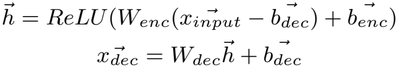

where *W*_*enc*_ is the SAE encoder weight matrix, *W*_*dec*_ is the SAE decoder weight matrix, 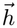 is the sparse hidden activation vector, 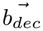 is the SAE encoder bias vector, and 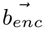 is the SAE decoder bias vector. The loss function combines the mean difference between the dense inputs and outputs of the SAE (MSE Loss) with the mean absolute activations of the sparse hidden activations (Sparse Loss). The loss function is given by:

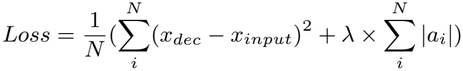

where *N* is the number of inputs, *x*_*dec*_ is the decoded output vector from the SAE, *x*_*input*_ is an activation vector from a ProteinMPNN encoder layer, *λ* is the sparse loss weight, and *a*_*i*_ is one activation value n in the SAE.

The PyTorch Paszke et al. (2019) Adam optimizer was used for training. The default beta parameter was used and a learning rate of 1e−4 was selected based on work by Bricken et al. (2023). During training, some dimensions in L1 SAEs show no activation for the last 5,000 batches. In this event, corresponding weights are reinitialized with a Kaiming uniform distribution Bricken et al. (2023).

During the course of training, several criteria were considered to predict how interpretable the model was before the computationally expensive evaluation procedure. Models were evaluated based on training and validation sparse and MSE loss, the number of dead neurons for the last epoch, the number of dead neurons since the last reinitialization Bricken et al. (2023), the percentage of non-zero activations (termed specificity), and model accuracy. Accuracy was calculated by feeding all three layers of SAE dense encodings back through the model during validation. An ideal interpretable model should have a low specificity value (each dimension only activates for a few samples and thus relates to very specific features) while having zero dead neurons (each neuron responds to inputs somewhat frequently) while maintaining high accuracy.

### 2.2. SAE Evaluation

#### 2.2.1. Feature Dataset

Models are interpretable if a human can trace its reasoning and understand the cause of a prediction Miller (2019). Based on work in Large Language Models (LLMs), models are known to optimize learning and storing features about the input that are then used to create predictions. Knowing which features are stored can be used to justify predictions and eventually establish a causal relationship. To show this for ProteinMPNN, sparse activations can be correlated to the presence of per-residue features where significant correlations mean a dimension is highly likely to store information for the given feature.

The per-residue feature dataset was created using the NCBI protein database Altschul et al. (1990). A search for X-ray diffraction structures with a resolution less than 2.5 Å, less than 500 residues, and less than 50% sequence identity produced 100,000 proteins with 25,000 representative structures. Of these representatives, 1,200 were randomly selected and their structures were downloaded from the RCSB PDB database Berman et al. (2000). 51 files were discarded for having incomplete or corrupted structures leaving a total of 1,149 structures. For each structure, per-residue features were found or calculated using MDAnalysis and Biopython Michaud-Agrawal et al. (2011); Gowers et al. (2016); Cock et al. (2009) giving a total of ~430,000 samples. Features are summarized in Table 1. The selected features were chosen based on their perceived significance to protein folding. Additionally, XY/XZ/YZ frame orientation angles were chosen since they were cited as an input in the ProteinMPNN paper but were not actually used since they provided no additional accuracy. It would be interesting if there were no improvement in accuracy since these angles were uncovered by the model. B-factor was selected since ProteinMPNN trained off of X-ray crystallography structures and may have learned some uncertainty factor proportional to the B-factor. Polarity categorizations were based on Trinquier and Sanejouand (1998) and volume categorizations were based on Darby and Creighton (1993).

**Table 1.** Residue-level features for evaluation.

| Feature | Description |
|---|---|
| **B-factor** | C<sub>α</sub> B-factor obtained from the PDB |
| **XY/XZ/YZ angles** | Angles describing residue frame orientation, computed from the three C<sub>α</sub> atoms defining the local C<sub>α</sub>–C<sub>α</sub>–C<sub>α</sub> plane. |
| **Close contacts** | Number of atom-atom salt bridge, hydrophilic, and hydrophobic contacts within 4, 3.5, and 2.5-4.6 *Å* respectively. Calculated with MDAnalysis |
| **Distance contacts** | Number of atom-atom pairs within a certain distance (16 bins spanning 2 to 22 *Å*). |
| **AA function** | Amino acids grouped by chemistry (Hydrophobic: GAVILM, Hydrophilic: CPSTNQ, Positive: KRH, Negative: DE, Aromatic: FYW) |
| **AA polarity** | Amino acids grouped by polarity (Hydrophobic: WCMIFLV, Medium: GRSTAP, Hydrophilic: EDKNQHY) |
| **AA shape** | Grouped into structural categories (Small: GA, Short chain: SCVT, Medium with branch: DNIL, Long with branched end: EQ, Long with no branch: KM, Very long with branched end: R, One ring: PHFY, Two rings: W) |
| **AA volume** | Grouped by Van der Waals volume of side chain in *Å*<sup>3</sup> (48: G, 67: A, 73: S, 86-96: CPDTN, 105-118: VEQH, 124: ILM, 135-148: KFYR, 163: W) |
| **Secondary structure** | Residue secondary structure assigned using implementation of DSSP in BioPython. |
| **SASA** | Solvent accessible surface area calculated with BioPython. |
| *ϕ*/*ψ* **angles** | Backbone dihedral angles calculated with BioPython. |

All angles were normalized to be between 0 and 180. The same 1,149 protein dataset was used as input for ProteinMPNN with the trained SAE and the sparse activations for all SAEs for each residue were collected.

Features were compared to SAE activations after each dataset was cleaned for duplicate entries or NaN values. Categorical comparisons were evaluated with ROC (receiver operating curve) AUC (area under the curve) and quantitative comparisons were evaluated with a Pearson test using the SciPy Python Library Virtanen et al. (2020). Categorical comparisons required at least 40 samples from each group and were downsampled to balance each group. Significant correlations were those with an ROC AUC value above 0.7 or a Pearson r correlation coefficient above 0.5. These thresholds were chosen as a generous lower bound to capture all potential correlations. Pearson correlations were only reported for those with a p-value less than 0.05 divided by the number of dimensions being tested against (i.e. 0.0002 for SAE256) to eliminate p-hacking bias.

#### 2.2.2. Sparse Embedding Clustering

With each message passing step, additional information changes how the latent space is optimally organized. These changes can be visualized via t-distributed stochastic neighbor embedding (t-SNE) van der Maaten and Hinton (2008) which reduces the number of dimensions in each vector to two. t-SNE was chosen since its reduction method preserves local distance between points. Evaluations in a 2D space cannot provide quantitative evidence of features since an interpretable model should have many unique, orthogonal dimensions. Rather, t-SNE provided a qualitative metric to observe how the distance between structurally dissimilar residues evolves.

Encodings for each layer were randomly downsampled to 1% of their original size, standardized, and labeled based on the secondary structure of the residue. The model’s ability to discern the absence of features as a negative control was also tested. Since it is hypothesized all features are based on structure and intrinsically disordered proteins (IDP) contain nearly none of these features, they may occupy a unique position in the latent space. If not, the latent space may not be ordered principally by structure. For this purpose, four IDPs (Q6UNL8, P27058, P04984, and Q2GI62) were selected from DisProt Nugnes et al. (2026) based on their high “disorder content” and “AlphaFold very low confidence content”. Their encodings were captured by the chosen model and displayed using t-SNE reduction.

#### 2.2.3. Visualizing Residue Activations

Previous attempts at protein design interpretability Simon and Zou (2025) have used a combination of the previously described quantitative features as well as more specific features from the UniProt Database Consortium (2022). However, these databases, while extensive, unfortunately fail to completely annotate all the known features in our evaluation dataset. SAE models may store information as features that are not used by humans such as distinguishing between the N-terminus of C-terminus of an *α*-helix, the presence of either of two or more seemingly unrelated amino acids, or a particular level of torsional strain.

For LLMs, SAE analyses have used manual annotation to describe the tokens (words or word segments) that activated a dimension the most and their confidence in that description Bricken et al. (2023). A similar procedure was employed to analyze each dimension of the third layer of SAE256. The residues that led to the greatest activation of each dimension were displayed and individuals saw the residues highlighted in the context of the full protein alongside secondary structure, amino acid identity, SASA, and any UniProt annotations (UniProt annotations are collected with the help of software provided by Galardini and Wagih (2020)). Visualizations were interactive and individuals annotated the similarities and their confidence in their description. High confidence features across multiple users were reported. Features were not cross-validated with a large number of users and qualitative data is subjective so no significance data is reported and not definitive proof of the existence of features.

#### 2.2.4. Steering

To move from justification to explanation, a causal relationship between the information in the latent space and predictions must be established Biran and Cotton (2017). If such a causal relationship exists, manipulating the sparse latent space would enhance or reduce the presence of protein features in the downstream protein sequence and folded protein. To prove this, the last layer of the chosen SAE was used to increase the thermostability of an scFv.

Selected sequences were compared against the original structure, a sequence generated by ProteinMPNN without an SAE, and a sequence generated by GeoEvoBuilder Liu et al. (2025) which is designed to improve thermostability without affecting structure.

The template structure Brolucizumab, an FDA approved scFv for neovascular age-related macular degeneration, was chosen. The sequence was obtained from the international Immunogenetics information system Lefranc et al. (2009). The native structure was obtained by folding this sequence with AlphaFold3 Abramson et al. (2024) with default settings. GeoEvoBuilder was used to generate 1000 sequences with default settings and the top scoring sequence was selected. ProteinMPNN was used to generate 1000 sequences and a consensus sequence of the most common amino acid at each position was selected.

Previous studies have increased the thermostability of the Fv fragment de novo by increasing non-covalent interactions in the V_H_/V_L_ interface in Tan et al. (1998); Ewert et al. (2003); Chowdhury and Vasmatzis (2003). Preventing unfolding of the chain limits aggregation and maintains functionality. To increase thermostability using SAE steering, four dimensions in the last layer of the SAE that best correlate with small branched chain amino acids (BCAAs) were identified. The values at these dimensions were increased for residues with less than 20% exposed surface area as determined by MDAnalysis. Altered sparse activations were fed into the SAE decoder and the ProteinMPNN decoder to obtain candidate sequences. Four candidate sequences were generated corresponding to the value added to their activation: mod=0, mod=0.5, mod=1, and mod=2. Candidate and baseline sequences were folded using AlphaFold3 with default settings and the first model was selected.

#### 2.2.5. Molecular Dynamics Simulations

To quantitatively assess thermostability, molecular dynamics simulations were performed. Proteins were solvated and ionized using VMD Humphrey et al. (1996) in a 70 Å cube with a 0.15 M potassium chloride solution. The minimized proteins were then equilibrated to the target temperature using NAMD Phillips et al. (2005). Each protein was simulated for 5 independent runs at 298 K, 373 K, and 433 K for 100 nanoseconds with 1 picosecond intervals (100,000 frames) using CHARMM36m forcefields Huang et al. (2017). All simulations were conducted using the Santa Clara University WAVE High Performance Computing cluster, with each run taking roughly 22 hours using an NVIDIA A16 GPU.

The simulation data was then processed using MDAnalysis. The root mean square deviation (RMSD) between alpha carbons of the first and current frame was calculated to quantify protein unfolding. Residues 111 to 133 were excluded since they are an unstructured region connecting the two chains, and their fluctuations are not indicative of overall stability. RMSD values at or below 2 Å correspond to a stable structure with little flexibility. Values above 2 Å correspond to large deviations from the starting position, indicating a degradation of the secondary structure. The mean fluctuation for each residue’s alpha carbon (RMSF) at 433 K was calculated using MDAnalysis and was used to compare the thermostability of each structure.

## 3. Results

### 3.1. Model Training

Each SAE three-layer model run used a batch size of 10,000 tokens (residues), 10,000 proteins per epoch, a learning rate of 0.0001, a sparse weight of 0.00025, dead neurons were reinitialized every 5,000 steps, and trained for 150 epochs. During initial testing, only the sparse weight was optimized. The learning rate was kept low and the SAEs were trained after loss had plateaued to encourage additional feature learning. The ideal sparse weight was found by decreasing it until the number of dead neurons started to increase. Well performing models after evaluation had higher final sparse loss values than expected. Loss values reported in Table 2 are averaged across all three layers while specificity and dead neurons are for Layer 3.

**Table 2.** SAE Model Final Epoch Validation Values.

| Models | SAE128 | SAE256 | SAE512 | SAE1024 |
|---|---|---|---|---|
| Sparse Loss | 3220.925 | 1736.604 | 1072.814 | 1317.327 |
| MSE Loss | 0.061 | 0.035 | 0.023 | 0.028 |
| Specificity | 0.990 | 0.778 | 0.649 | 0.548 |
| Dead Neurons | 0 | 3 | 2 | 0 |

### 3.2. Feature Correlation

Each model was used to create a set of encodings of the evaluation dataset. These encodings were then randomly downsampled to 10% of their original size (to 43,000 samples) and correlated to the evaluation dataset features. The number of dimensions that were well correlated to each feature is reported in Fig. 2. Some dimensions correlate well to multiple features such as the presence of an *α*-helix and the absence of an unstructured region. The high number of dimensions correlated with secondary structure and side chain volume indicates the model performed best when much of the latent space was used to store this information and recovering these features was important for sequence prediction. Correlations may or may not be causal, and the actual relationship can only be determined with future experimentation such as prediction steering. As mentioned previously, the final encoding should be the most information-rich and contain the most features. The greatest correlation value for every SAE Layer 3 dimension for some features is listed in Table 3. AUC or r values may represent the inverse feature (i.e. absence of an alpha helix).

**Table 3.** Greatest Correlations for SAE Layer 3 Models (Highest values in bold)

| SAE Model | None | 128 | 256 | 512 | 1024 |
|---|---|---|---|---|---|
| No Structure | 0.639 | 0.744 | **0.802** | 0.750 | 0.767 |
| Beta Strand | 0.650 | 0.786 | **0.826** | 0.768 | 0.762 |
| Alpha Helix | 0.744 | 0.747 | 0.778 | **0.793** | **0.793** |
| Function | 0.664 | 0.748 | **0.795** | 0.789 | 0.736 |
| Polarity | 0.725 | 0.727 | 0.801 | **0.819** | 0.713 |
| Shape | 0.654 | 0.759 | **0.779** | 0.752 | 0.765 |
| Volume | 0.694 | 0.894 | 0.893 | **0.912** | 0.897 |
| SASA | **0.766** | 0.521 | 0.581 | 0.494 | 0.516 |
| Phi/Psi Angle | **0.767** | 0.469 | 0.467 | 0.589 | 0.510 |

**Figure 2.**
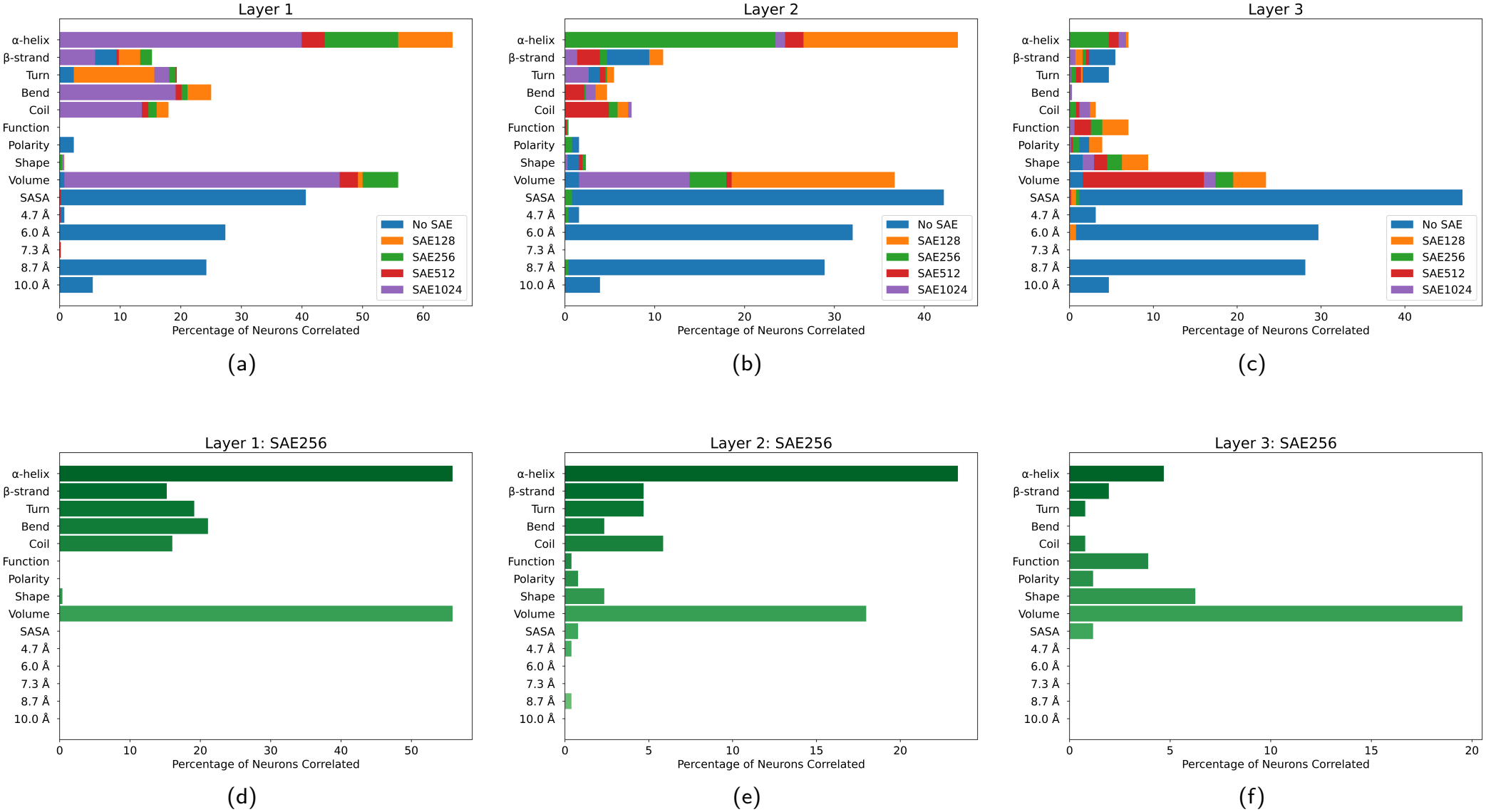
Percentage of dimensions correlated with features for all SAE models. (a, b, c) Every SAE model was compared against no model for layers 1-3. The distribution is consistent across all models but changes at different layers. (d, e, f) The number of correlated dimensions of SAE256 was reported for all three layers. The number of correlated dimensions decreased with successive layers.

SAE256 and SAE512 performed the best. The first two layers for each reported a large percentage of dimensions dedicated to structural characteristics like secondary structure and phi and psi angles. The third layer for each showed a number of neurons related to chemical features such as amino acid function, shape, and SASA. These features informed the final amino acid identity prediction. As the model progresses, each layer learned more information about the protein and obtained a less structural and more functional understanding.

However, higher expansion levels (8 times larger) reported a similar distribution of well correlated neurons and does not uncover features such as the number of contacts or greater strength correlations (Table 3). Many features were not uncovered by any of the models including the XY, XZ, or YZ frame orientation, B-factor, or the number of hydrophobic, hydrophilic, or salt-bridge contacts.

An SAE model with no expansion was similarly tested and had the same distribution of features as other SAE models (Fig. 2).

The original ProteinMPNN encodings were also evaluated, and many significant correlations were found for beta strands, polarity, exposed surface area, and the number of contacts at certain distances (Fig. 2). However, the number of correlated neurons between layers did not change, and the distribution was very different. Further analysis of these correlations showed that these features were unrelated to sampling bias or other factors. Since these features were related to the number of contacts a residue has, these correlations may be a remnant of the initial embedding where many contacts would give a large activation value before any message passing steps. However, the reasoning is still unclear and requires further study. SAE models of any size were able to uncover features that are not correlated in the original model, including the presence of an alpha helix, beta-bridge, 3-10 helix, turn, absence of secondary structure, phi and psi angles, and increase the proportion of the latent space related to turns, residue shape, and residue volume. While this information was stored in the dense latent space, it could not be revealed or understood without SAEs. Using the correlation data, SAE256 was chosen for further analysis. This model captures as many features as larger models and since it is smaller, its behavior can be changed by manipulating only a few dimensions.

### 3.3. Clustering

To see how the latent space of SAE256 is organized, and how that organization changes through the model, the encodings were displayed with t-SNE. The number of encodings is randomly downsampled to 4,304, and the t-SNE reduced encodings are shown in Fig. 3. The scatterplot for the first layer showed a distinct separation of clusters based largely on secondary structure, while the second and third layer scatterplots had a more uniform distribution of secondary structure. The distance between samples of differing secondary structure likely decreased because other dimensions encapsulated new information that was mutually inclusive of secondary structure. These plots support the theory that later layers encapsulate new and non-structural information about the protein and is supported by the increase in dimensions well correlated with residue function, polarity, and shape in last layer of the SAEs.

**Figure 3.**
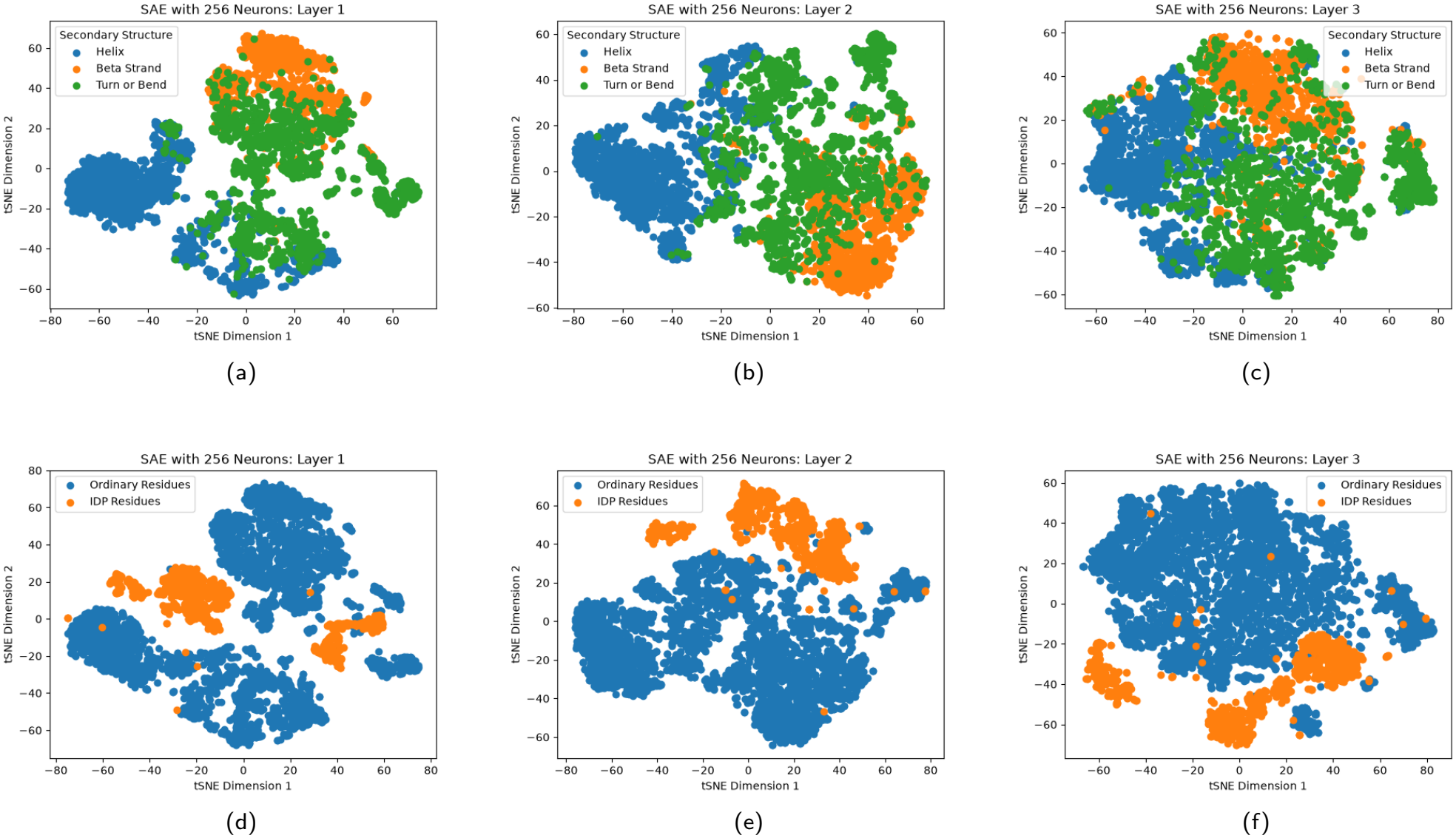
t-SNE visualization of SAE256 sparse encodings. (a, b, c) 4,304 256-dimension encodings for each of the three layers were reduced to 2 dimensions using t-SNE and were labeled based on their secondary structure. The same encodings were shown in blue alongside intrinsically disordered protein (IDP) residues in orange for all three layers (d, e, f). IDP residues formed clusters distinct from other residues in all three layers.

As for residues in IDPs, the encodings formed a distinct cluster away from residues of any other proteins. These plots provide evidence that while secondary structure alone was not the sole indicator of sample position within the sparse embeddings, the organization of the sparse latent space was still largely governed by structure. If this was not the case, IDP encodings would be uniformly distributed across the latent space similar to the turn or bend secondary structures.

### 3.4. Qualitative Feature Detection

Each dimension for the third layer of SAE256 further examined by assessing the top 10 residues that lead to the greatest activation. Several features were identified and images of the corresponding residues are in Fig. 4. Disulfide bonds, coordination to zinc atoms, and coordination or binding to heme or other molecules were present in the majority of the top activations for some dimensions. Dimensions 1 and 116 are shown in Fig. 4 where the residues that most strongly activate these dimensions are primarily cysteines bound to heme molecules and zinc ions respectively.

**Figure 4.**
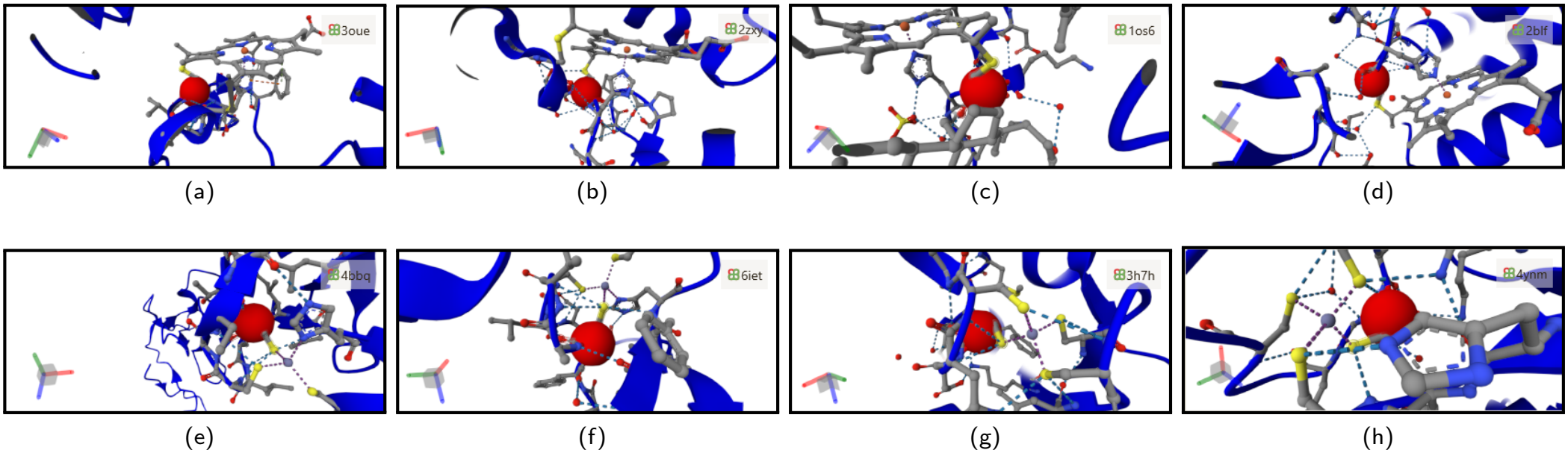
Most active residues for SAE256 Layer 3 Dimensions. Of 430,000 residues, the 1st, 6th, 7th, and 8th (a, b, c, d) greatest activations in dimension 1 all were cysteines bound to a heme molecule. The 2nd, 3rd, and 4th greatest activations were of cysteines bound to heme in the same protein as (a). The 1st, 4th, 5th and 6th (e, f, g, h) greatest activations in dimension 116 all were cysteines bound to a zinc ion. 3rd greatest activation was for the same protein as (e). The 2nd greatest activation was a disulfide bridge.

### 3.5. SAE Model Steering

scFvs have previously been thermostabilized by increasing the hydrophobicity of the interior pocket. To do so for Brolucizumab, the dimensions in the third layer of SAE256 related to branched chain amino acids (BCAAs) (leucine, isoleucine, and valine) were found: 46, 48, 182, 191, and 221 (Table 4). Using a 20% SASA threshold, 104 buried residues out of the total 252 residues were changed. At these buried residues, sparse activations of the chosen dimensions were increased by 0 (control), 0.5, 1, and 2.

**Table 4.** Pearson *r* Correlation Coefficient Values for SAE256 Layer 3 Dimensions of BCAAs.

| Dimensions | 46 | 48 | 182 | 191 | 221 |
|---|---|---|---|---|---|
| Leucine | 0.723 | 0.791 | 0.624 | 0.676 | 0.741 |
| Isoleucine | 0.493 | 0.595 | 0.849 | 0.765 | 0.795 |
| Valine | 0.409 | 0.454 | 0.838 | 0.761 | 0.784 |

The produced candidate sequences were folded with AlphaFold3 Abramson et al. (2024). Their structures were aligned using Berman et al. (2000) (Fig. 5). RMSD analysis of MD simulations (Fig. 6) showed no substantial unfolding for any proteins at 298 K. Baseline proteins of the original sequence, ProteinMPNN seqeunce, and GeoEvoBuilder sequence and mod=2 exhibited substantial unfolding at 433 K, while mod=0, mod=0.5, and mod=1 showed lower RMSD values throughout the simulation. For these candidates RMSD values leveled off around 5 Å, indicating an alternate stable conformation. While mod=0.5 and mod=1 showed marked improvement in thermostability over the baseline structures, so did mod=0. These findings are consistent with the mean RMSF which averages all C*α* atom fluctuations across time for each protein at 433 K (Fig. 7). SAEs introduce some error into the latent space which is minimized during training (MSE error). The final MSE loss may have shifted predictions in favor of more hydrophobic amino acids meaning the presence of the SAE model itself has introduced some thermostability.

**Figure 5.**
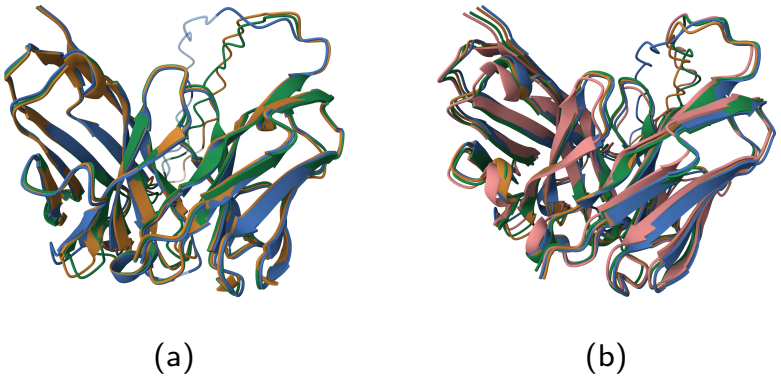
Structures aligned with RCSB PDB Pairwise Structure Alignment with TM-align. (a) Native scFv in orange, ProteinMPNN variant in blue, and GeoEvo variant in green. (b) mod=0 in orange, mod=0.5 in blue, mod=1 in green, and mod=2 in pink.

**Figure 6.**
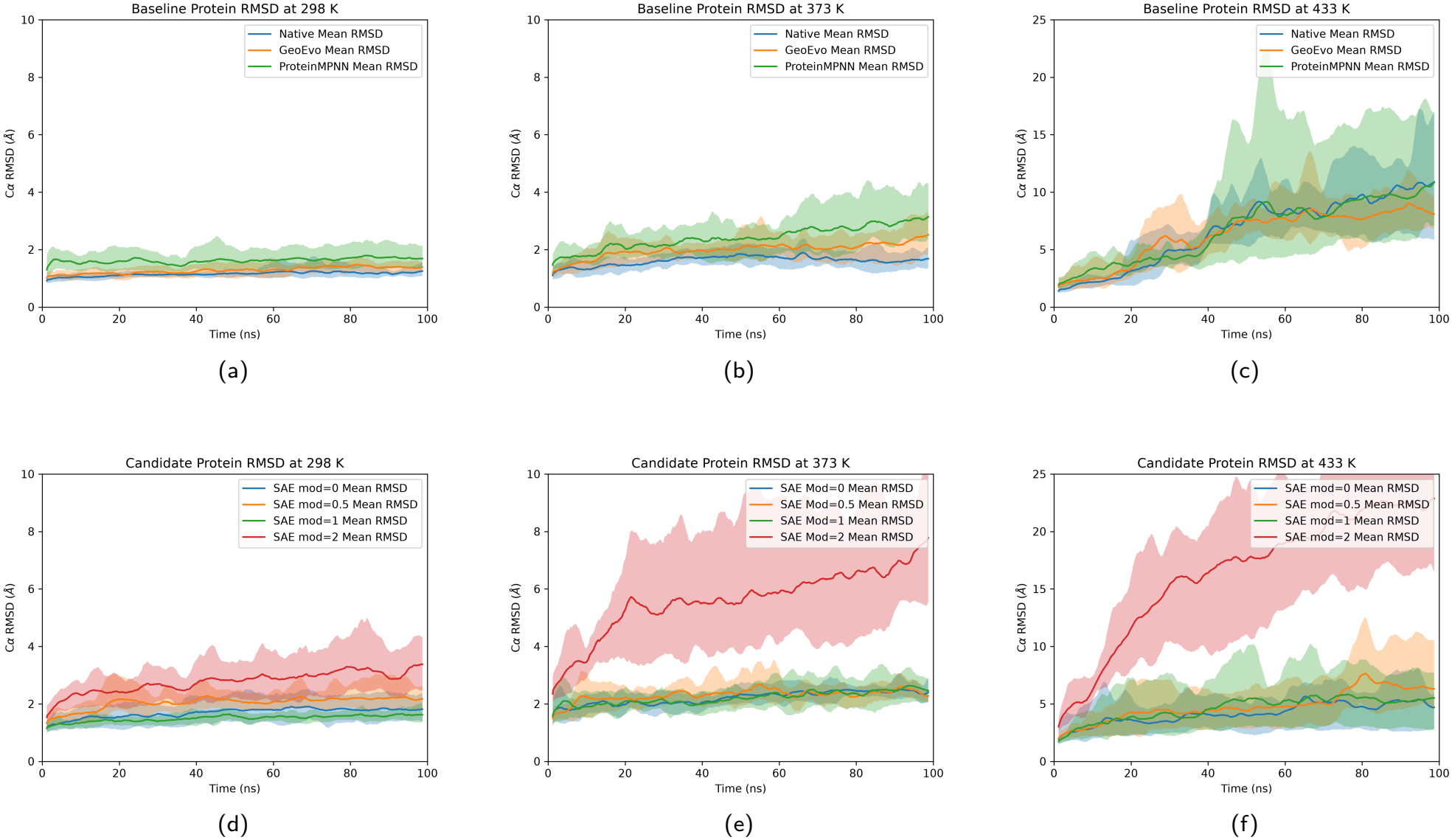
RMSD for baseline and candidate proteins. RMSD for for the native structure, ProteinMPNN variant, and GeoEvo variant (a, b, c) and four SAE modified structures (d, e, f) are shown in band plots. The dark line is the mean across all 5 trials and the band is bound by the minimum and maximum RMSD. Values are smoothed with rolling average window of 25 frames. Plots show significant unfolding for all baseline structures at 433 K and only moderate flexibility for candidate structures except for SAE mod=2 which unfolds at 373 K and 433 K.

**Figure 7.**
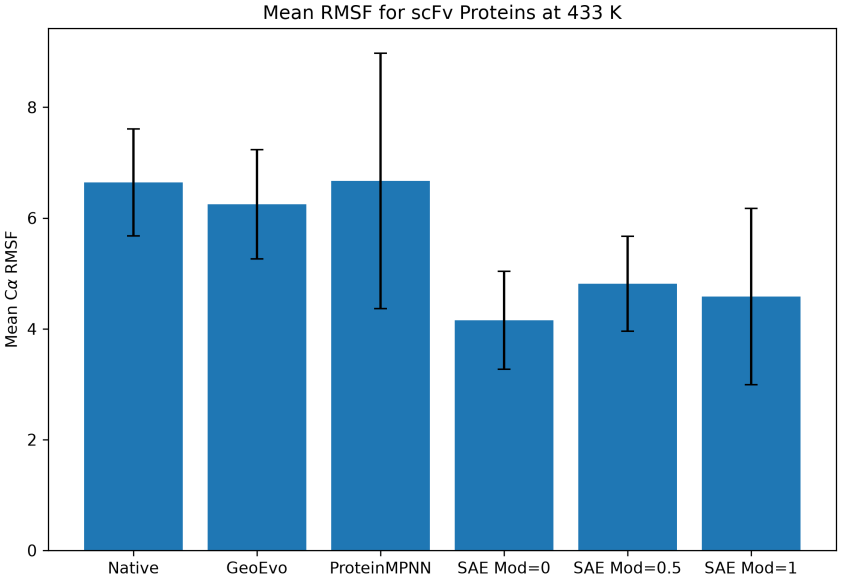
Average residue RMSF at 433 K with error bars for 95% confidence interval.

To validate this claim, a smaller protein, egg white cystatin (PDB ID: 1CEW), was selected to undergo the same steering procedure and only dimension 221 (correlated to BCAAs) was changed. For each change in activation, 1000 sequences were generated, and the proportion of nonpolar (WCMF), branched chain (ILV), intermediate (GRSTAP), and polar (EDKNQHY) amino acids was determined. (Table 5).

**Table 5.**
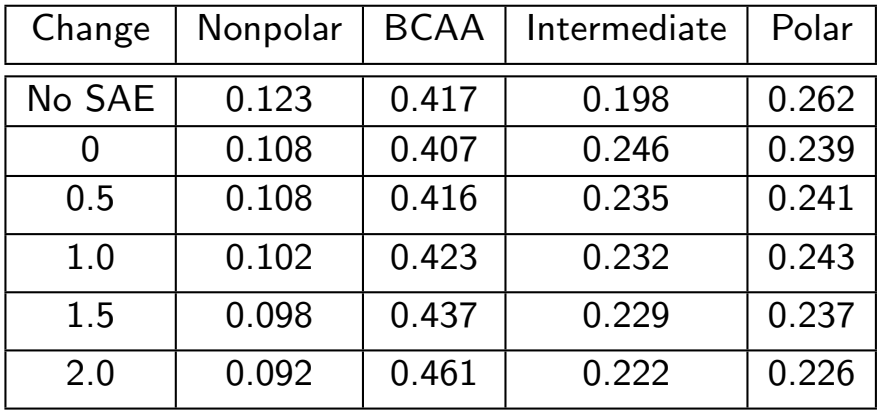
Proportion of Buried Amino Acid Type for Change in Sparse Activation.

| Change | Nonpolar | BCAA | Intermediate | Polar |
|---|---|---|---|---|
| No SAE | 0.123 | 0.417 | 0.198 | 0.262 |
| 0 | 0.108 | 0.407 | 0.246 | 0.239 |
| 0.5 | 0.108 | 0.416 | 0.235 | 0.241 |
| 1.0 | 0.102 | 0.423 | 0.232 | 0.243 |
| 1.5 | 0.098 | 0.437 | 0.229 | 0.237 |
| 2.0 | 0.092 | 0.461 | 0.222 | 0.226 |

There was a decrease in the proportion of hydrophobic and BCAA amino acids in buried positions after the introduction of the SAE which slowly increases with greater activation of dimensions 221. Interactions between dimensions may also be hampering the effectiveness of steering. The bias of each SAE model was evaluated using a similar procedure. With 1CEW, no activations were changed but 1000 sequences were generated with each model and the proportion of each amino acid type was measured and shown in Table 6.

**Table 6.** Proportion of Buried Amino Acid Type With No Modification.

| Model | Nonpolar | BCAA | Polar | Other | MSE Loss |
|---|---|---|---|---|---|
| No SAE | 0.123 | 0.417 | 0.262 | 0.198 | NA |
| SAE128 | 0.123 | 0.415 | 0.262 | 0.200 | 0.061 |
| SAE256 | 0.108 | 0.407 | 0.239 | 0.246 | 0.035 |
| SAE512 | 0.111 | 0.498 | 0.215 | 0.176 | 0.023 |
| SAE1024 | 0.070 | 0.433 | 0.197 | 0.301 | 0.028 |

Comparing each proportion to the corresponding value without an SAE, each model changed the amino acid distribution irrespective of the magnitude of MSE loss. Indeed, SAE256 decreased the proportion of polar amino acids while SAE512 greatly increased the proportion of BCAAs. Models with more dimensions drifted further from the original distribution likely because small errors in the SAE compound as dimensions interact. So while the magnitude of MSE loss is not an indicator of performance, models trained for many more epochs, with more optimized hyperparameters, or with advances to training methodology are likely to perform better. The effect of this error is not related to the previous correlation analysis and is only relevant to the re-encoding step during steering.

## 4. Discussion

In summary, SAE models of any size recognize secondary structures, side chain volume, and residue function from the information in the ProteinMPNN latent space (Fig. 2). Additionally, SAEs are able to reconstruct non-trivial features such as coordination to zinc atoms or heme (Fig. 4). Since only some cysteine residues coordinate with zinc or heme, the model must recognize the coordination before the amino acid identity and use it to inform its prediction, not the other way around. These findings provide a great deal of mechanistic understanding to how ProteinMPNN works: it does implicitly recognize certain functional and structural per-residue characteristics and use these to inform amino acid prediction.

This is line with findings on the use of SAEs in LLMs (Bricken et al. (2023); Bills et al. (2023); Wang et al. (2022)) and image classification models (Liu et al. (2018); Zeiler and Fergus (2014)). In LLMs, segments of the sparse model stored information for features such as punctuation, locations, language, etc. (Bricken et al. (2023)). Image models contain dimensions storing information for curves, edges, or other motifs (Zeiler and Fergus (2014)). These features are used to form circuits that led to predictions (Cunningham et al. (2024); Wang et al. (2022) which may similarly be found in the ProteinMPNN decoder.

Automating feature analysis is still a large hurdle for SAE analysis. While larger models could uncover more protein features, examining every dimension manually with the current methods would be infeasible. Additionally, any method would need to be human-agnostic since models likely use features which do not seem natural to us. For SAE analysis of LLMs, Bills et al. (2023) used language models to generate descriptions of the tokens for top activations and predict the activation of a sample token based on that description. Adapting this to protein design, future methods could perform a local structure alignment on top residues similar to Simon and Zou (2025) and compare that environment to that of a test residue to predict the sparse activation of the test residue based on their similarity. Such a method would account for any and all potential features or motifs.

Experiments with using SAEs to steer predictions are also promising even though they require much more study. The procedure developed in this paper could be used for any protein with a large interior volume. Much like how SAE steering on LLMs could change word meanings Bricken et al. (2023); Cunningham et al. (2024); Wang et al. (2022), SAE steering on

ProteinMPNN could in the future be used to insert features such as zinc finger motifs, disulfide bridges, or secondary structures. Such a methodology has advantages over de novo design since the model always works to compromise the user’s alterations with its understanding to create the most stable sequence and structure. Although, some alterations are so significant they create a very unstable protein as shown by mod=2 (Fig. 6).

However, this procedure is currently limited by our understanding of the mechanics of the ProteinMPNN decoder and the precision of sparse dimensions. Future steering attempts should reduce MSE loss more during SAE training to limit the influence of the SAE on node encodings. More training could also allow the model more time to learn more specific and uncommon features. Combined with thorough qualitative analysis, steering could be an incredibly useful and multi-purpose tool for protein engineering. Still, further work is needed to more completely understand ProteinMPNN’s mechanics before the ultimate goal of fully interpretable and custom protein design.

## 6. Acknowledgments

The authors thank the anonymous reviewers for their valuable suggestions as well as Santa Clara University for access to the HPC WAVE Supercomputer. This work is supported in part by Santa Clara University via the ALZA and 2FURS grants. The authors would also like to thank Liam Pruitt and the other members of the McCully Lab.

## 7. Data availability

The source code for this article is available at github.com/neelmukkavilli/SAEProteinMPNN.git

## 8. Author contributions statement

N.M. conceived the procedure for the architecture, training, correlation, and MD simulations. N.M. and K.I conducted training, ran the experiments, and analyzed the results. N.M., M.E.M., D.A.C.B., and H.A. conceived the procedure for clustering and steering. N.M. and K.I. conceived the procedure for qualitative feature discovery. N.M. and M.E.M. analyzed the MD simulations. K.I. developed a website for qualitative feature analysis. N.M. wrote the manuscript which was reviewed by K.I., H.A., M.E.M., D.A.C.B., and D.A..

